# Glucose levels impact the morphology and cell-type composition of human cerebral organoids

**DOI:** 10.1101/2025.10.02.680109

**Authors:** Gautami R. Kelkar, Balaji M. Rao, Albert J. Keung

**Affiliations:** Department of Chemical and Biomolecular Engineering, North Carolina State University, Raleigh, NC, USA; Golden LEAF Biomanufacturing Training and Education Center, North Carolina State University, Raleigh, NC, USA

**Keywords:** Cerebral Organoids, Glucose concentration, Cell culture media, Morphology, Cell-type composition, Neuronal differentiation, Metabolism

## Abstract

Human cerebral organoids, derived from pluripotent stem cells, are powerful models for studying human brain development. The understanding of how morphogens can be used to guide patterning and differentiation has matured rapidly, however, the influence of basal media components on organoid development remains unclear. Standard organoid media frequently contain non-physiological concentrations of nutrients, including glucose, a central regulator of cellular metabolism and signaling. Here, we examine how glucose availability shapes cerebral organoid growth, morphology, and cell-type composition by comparing conventional hyperglycemic media to media with glucose levels more closely resembling physiological conditions. We find that organoids derived from multiple human pluripotent stem cell lines can grow in physiological glucose, but exhibit altered growth rates, structural features, and lineage distributions. In H9 embryonic stem cell-derived organoids, inhibition of the mTOR pathway under physiological glucose restores neurodevelopmental cell types otherwise diminished in these conditions. These findings highlight glucose as a key determinant of organoid lineage specification and cellular signaling. Importantly, however, glucose modulation does not reduce variability across organoids or cell lines, underscoring the need to better understand and control sources of heterogeneity to improve organoid models.

## 1. Introduction

Human cerebral organoids (hCOs) are derived from embryonic (ESC) or induced pluripotent stem cells (iPSC) and can recapitulate some of the complex architectures of the human tissue *in vivo* (1,2). As they provide a human-specific platform for *in vitro* studies, there is a strong motivation to continue enhancing their capabilities. Media formulations in particular have profound effects on hCO derivation. A major focus in the field has been identifying combinations of morphogens to mimic and capture different regions and cell types of the brain (3–12).

Basal media components also govern cell differentiation, but their role is understudied. These components are largely derived from traditional formulations created decades ago for immortalized cell lines; as such, media used for hCO generation have non-physiological concentrations of several media components, particularly glucose. Studies comparing hCOs grown *ex vivo* to those transplanted into a mouse brain hint at the non-physiological aspects of culture media altering cell type composition and contributing to low fidelity and reproducibility in hCO cultures (13). Similarly, a previous study of 3D embryoid bodies suggests that glucose concentration in the media affects growth and morphology (14). Indeed, studies in 2D cultures have found that modifying nutrient concentrations in the media to match physiological levels might improve neuronal differentiation, maturation and function (15,16).

We focus here on the potential role of glucose since it is the primary source of carbon and energy (17–19). Moreover, it interacts with other metabolic and cell signaling pathways to contribute to cell fate decisions (20–24). hCOs have been used to study the role of glucose in disease states such as diabetes (25) and ischemia (26). However, its role during standard hCO development is often overlooked. Importantly, typical hCO protocols supply hyperglycemic quantities of glucose to pluripotent stem cells (14,27) and raise practical questions about the physiological relevance of conclusions drawn from these conditions and if current conditions should be modified.

In this study, we investigated the morphological and cell-type differences between standard high (13-17 mM) versus near physiological glucose (3.5 mM) conditions for hESC and iPSC-derived hCOs. It is important to note that, while the levels used in this study were chosen to reflect those expected under physiological conditions, the precise *in vivo* concentrations are not known (28). We demonstrate that physiological glucose conditions can support hCO growth for ∼10 days, although the growth rates, which correlate with glucose consumption patterns, are expectedly slower than high glucose conditions. Interestingly, even at physiological concentrations, glucose in the media does not appear to be fully consumed between media changes. Nevertheless, we observe that for multiple cell lines, physiological glucose leads to loss of neurodevelopmental cell types. However, for H9 ESC-derived hCOs, the slower growth rate and lost cell types in physiological glucose conditions can be partially rescued by inhibiting the mTOR signaling pathway. Additionally, we find that each PSC line used in this study has a unique phenotypic response to glucose levels during hCO generation, drawing attention to the inherent variability of hCO models.

## 2. Materials and methods

### 2.1 Human Pluripotent Stem Cell (hPSC) culture

Eight hPSC lines (Supplementary table S1) were maintained on growth factor-reduced Matrigel (Corning, USA) coated 6-well plates (Corning COSTAR^™^, Fisher Scientific, USA) in mTeSR Plus (StemCell Technologies, Canada). Cells were passaged every 3-5 days as necessary using 0.5 mM EDTA (Invitrogen, USA).

### 2.2 hCO culture

Whole-brain hCOs were generated and maintained by modifying the protocol in (29). hPSCs were allowed to reach 80-90% confluency before they were dissociated into a single-cell suspension using Accutase (BioLegend, USA) and re-plated in low-adhesion U-bottom 96-well plates (Corning COSTAR^™^, Fisher Scientific, USA) at 10,000-13,000 cells/well. DMEM/F12 is used as the basal media for hCO generation. DMEM/F12 without glucose and l-glutamine (Biowest, USA) was first supplemented with 2.5 mM Glutamax (Gibco, USA). For high and physiological glucose conditions, glucose (Gibco, USA) was added to DMEM/F12 at 17.5 mM and 3.77 mM respectively. For the physiological glucose media, L-glucose (MilliporeSigma, Germany) was added to a concentration of 13.73 mM to control for osmolarity. For the hPSC growth stage (days 0-6) and the neural induction stage (days 6-11), the media consisted of ∼76% and ∼97% v/v DMEM/F12 respectively. The other media components for the hPSC growth stage include 19% KnockOut Serum Replacement (KOSR – Gibco, USA), 3% Fetal Bovine Serum (FBS – Avantor, USA), 1% GlutaMAX (Gibco, USA), 1% MEM-Non-Essential Amino Acids (Cytiva, USA), and 100 μM 2-mercaptoethanol (Amresco, USA), 50 μM Y-27632 (StemCell Technologies, Canada), and until day 4 - 4 ng/ml bFGF (Invitrogen, USA). Glucose concentrations for KOSR and FBS were measured using the Cedex^®^ Bio Analyzer (Roche Diagnostics, Switzerland) to calculate the final glucose concentration in the media. For the neural induction stage, the media also consists of 1% GlutaMAX, 1% MEM-Non-Essential Amino Acids, 1% N2 Supplement (Gibco, USA), and 1 μg/mL Heparin (Sigma Aldrich, Germany). Partial media changes were performed every two days, with a complete media change on day 6 to switch media types.

### 2.3 hCO size measurement

Images for size measurement were taken using an epifluorescence microscope with a documentation camera with a 4X objective (Nikon Instruments, Japan). Quantifications were performed manually in FIJI (30). The sizes of hCOs presented in each graph were normalized to the corresponding average size in physiological glucose at Day 2. The numbers of hCOs measured for each experiment can be found in Supplementary Table S2.

### 2.4 Glucose uptake measurement

hCOs were grown as described above. During media changes every other day, a pre-defined volume of media was removed from each well (75 μl on Day 2, 100 μl on Day 4, and 175 μl on Days 6 and 8) and spent media from 10 hCOs (8 for GM25256) were pooled into 1 sample. 3 such samples were collected for each condition, time point, and cell line. Samples were centrifuged at 300 g for 5 mins to remove cell debris, and identical volumes of supernatant were frozen at -80 ^○^C until analysis. Fresh media were similarly stored in triplicates and used as controls for the measurement. Glucose concentrations were measured on the Cedex^®^ Bio Analyzer (Roche Diagnostics, Switzerland) using the Glucose Bio kit. A pre-defined volume of fresh media was added to each well during each media change (150 μl on Days 2 and 4, 200 μl on Days 6 and 8). Volume fractions of spent and fresh media on each day were used to calculate the resulting glucose concentration after media changes i.e. the glucose availability on that day. Consumption rates are calculated on a per day basis.

### 2.5 Phenotype Characterization

Brightfield images of hCOs were captured at 4X magnification on an epifluorescence microscope with a documentation camera (Nikon Instruments) every other day from Day 2 to Day 10. hCOs were scored for 2 features - presence of an organized epithelial layer and cyst-like appearance. 3 levels of scores were used for each feature: 0 - complete absence of the feature, 0.5 - partially developed / visible feature, 1 – fully developed / clearly visible feature. All the hCO images were manually scored. For graphical representation, the epithelial layer was displayed on the X-axis and cyst-like features were displayed on the Y-axis. Each axis had 3 bins corresponding to the 3 scoring levels forming a 3 x 3 grid. Thus, each grid cell corresponds to one pair of coordinates or one type of hCO. Morphology distribution in both glucose conditions, at each time point was represented as a bubble plot, with the size of the bubble indicating the relative abundance of hCOs within a grid cell. Bubble sizes in each plot were scaled to the highest frequency within that plot. For each axis, +0.05 and -0.05 from the actual coordinates were used to stagger the scores to keep bubbles within the same grid cell visually separated. Relative positions of bubbles within each grid cell have no physical meaning. Comparison should be made across grid cells as a whole.

### 2.6 Histology and immunofluorescence

Tissues were fixed in 4% paraformaldehyde for 20 minutes at 4 ^○^C followed by washing in 1X PBS (Gibco, USA) three times for 10 minutes each. Tissues were allowed to equilibrate in 30% sucrose overnight and then embedded in 10% / 7.5% sucrose/gelatin. Embedded tissues were frozen in an isopentane and dry ice bath at -30 to -50 ^○^C and stored at -80 ^○^C. Prior to analysis, they were cryosectioned into 20-30 μm slices using a cryoStat (ThermoFisher, USA). For immunohistochemistry, hCO sections were blocked and permeabilized in 1% Triton X-100 and 5% normal donkey serum (Jackson Immunoresearch Labs, USA) in 1X PBS. hCO sections were then incubated with primary antibodies in 10% Triton X-100, 1% normal donkey serum in UltraPure^™^ water (Invitrogen, USA) and 10X PBS overnight at 4 ^○^C at the following dilutions: SOX2 (goat, R&D Systems, USA, 1:200), PAX6 (rabbit, abcam, UK, 1:100), Ki67 (rabbit, abcam, UK, 1:150), TUJ1 (mouse, MilliporeSigma, Germany, 1:500), and phospho-vimentin (mouse, MBL International, USA, 1:200). Following primary antibodies, sections were incubated with secondary antibodies - donkey Alexa Fluor 488, 546, and 647 conjugates (Invitrogen USA, 1:250) in 10% Triton X-100, 1% normal donkey serum in UltraPure^™^ water, and 10X PBS for 2 hours at room temperature, and the nuclei were stained with DAPI (Invitrogen, USA). Slides were mounted using ProLong^™^ Antifade Diamond (Invitrogen, USA). Images were taken with Nikon A1R confocal microscope (Nikon Instruments, Japan).

### 2.7 Image analysis and quantification

All images within a batch for each cell line were captured using identical microscope settings for each channel, and identically processed in FIJI (30) to create a maximum intensity z-projection, subtract background, and adjust channel intensities. A CellProfiler-automated pipeline was used to identify nuclei (DAPI) and quantify other cell-type markers as a fraction of DAPI counts in each image. For nuclear markers (SOX2, PAX6, and Ki67), DAPI+ regions that showed overlapping signal for these markers were selected. For cytoplasmic markers (TUJ1 and phosphor- vimentin), DAPI+ regions that were enclosed by signals for these markers were selected. Representative pipelines used in CellProfiler are available in the supplementary material. The values of parameters within the pipeline were independently selected to optimize quantification for each batch of a cell line. The number of hCOs measured in these experiments can be found in Supplementary Table S4.

### 2.8 Inhibitor treatment

H9 ESCs were chosen for inhibitor treatment due to their relatively greater reproducibility in high and physiological glucose. Rapamycin (MedChemExpress, USA) was reconstituted in DMSO (FisherScientific, USA) and was used at a final concentration of 20 nM in the media during hCO generation and replenished at every subsequent media change. Identical volumes of DMSO were added at each stage for vehicle control. Size measurement, phenotype characterization, and cell-type marker quantification were performed as mentioned above.

## 3. Results

### 3.1 Glucose affects hCO growth rate

To assess the impact of glucose levels on hCO growth, we observed whole-brain hCOs generated with the Lancaster protocol for 10 days (29). These hCOs are derived from 8 distinct hPSC cell lines. Two stages of early development are covered in this time frame – hPSC growth / embryoid body formation and neural induction. To mimic conditions close to those expected physiologically *in vivo*, we used 3.1-3.6 mM glucose in the media through 10 days (Figure 1A). This concentration range is based on the typical glucose concentrations that human embryos are exposed to *in utero* (31) and the levels used in *in vitro* embryo cultures (28). Up to 10 days in culture, we observed that physiological glucose conditions support hCO growth. It is important to note that the specific magnitudes of growth rates in both high and physiological glucose conditions are cell line and batch dependent (Figure 1B; Supplementary Figure S1). The sizes of hCOs in high vs physiological glucose conditions at day 4 were comparable. However, at 10 days, a majority of batches (∼75%) yielded smaller hCOs in physiological glucose conditions. Interestingly, while the final hCO sizes were smaller, the fraction of proliferative cells (marked by Ki67) in the hCOs at day 10 were significantly lower in only ∼25% of physiological glucose hCO cultures (Figure 1C).

**Figure 1.**
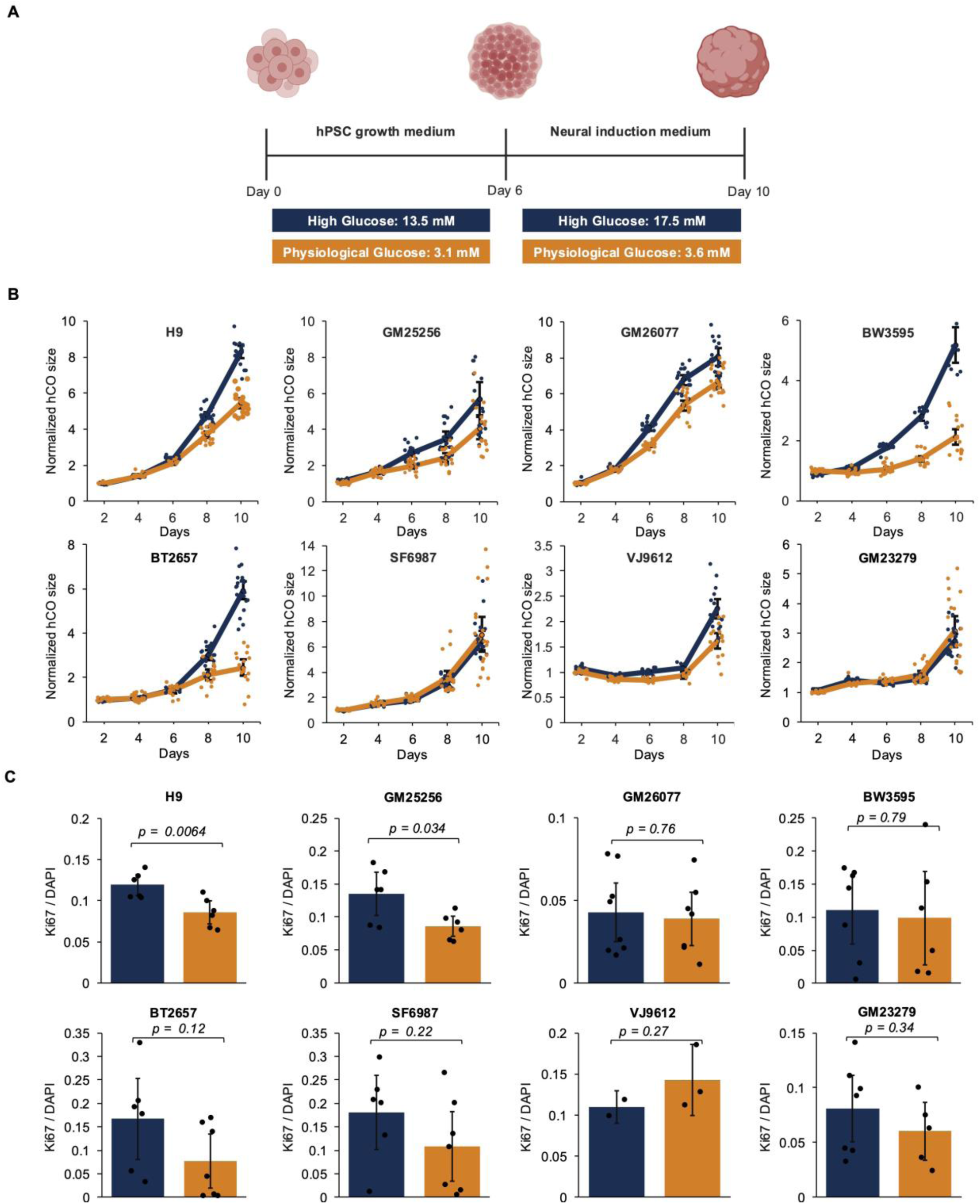
Glucose affects hCO growth rate. A) hCO growth protocol in high and physiological glucose conditions. B) hCO growth over 10 days in high (blue) and physiological glucose (orange) conditions for hCOs generated from different cell lines. hCO sizes on the y-axis are normalized to the mean size of physiological glucose hCOs on Day 2. Solid lines connect the mean hCO size between time points, dots represent individual hCO sizes, and error bars represent 95% confidence intervals. The numbers of hCOs measured can be found in Supplementary Table S2. C) Mean fraction of proliferating Ki67+ cells relative to DAPI (nuclei) for high (blue) vs physiological glucose (orange) conditions. Error bars represent 95% confidence intervals. Black dots represent individual hCOs used for immunofluorescence quantification. The numbers of hCOs measured are in Supplementary Table S4. Samples were compared using a two-sample t-test assuming unequal variances.

### 3.2 Glucose conditions and growth stage influence morphological heterogeneity in hCOs

Phenotypic variability is commonly observed within and between cell lines (32). Here we observed morphological diversity in hCOs derived from different cell lines along with diversity in glucose-induced effects. 11 types of hCO morphologies were observed over 10 days. Dimensionality reduction identified two dominant morphological features – epithelial layer and cyst formation (Supplementary Figure S2). hCOs were scored for these two features to enable quantitative comparison. The expectation based on standard hCO protocols is that with time, hCOs should start showing a prominent epithelium layer (29). However, we observe that 1 of the 8 cell lines (GM26077) show limited epithelial layer formation in typical cultures conditions, while 3 additional cell lines had a reduction in this phenotype in physiological glucose (BW3595, BT2657, GM23279) (Figures 2A, Supplementary Figure S3). In addition to the phenotypes themselves, we explored whether the magnitude of the variabilities in morphologies exhibit noticeable differences between conditions. The variabilities are represented by the standard deviation of the phenotype scores (Supplementary Figures S4 and S5). Across time, cell lines, and features, the variability between high and physiological glucose conditions is comparable (Figure 2B). Comparison between the two features demonstrates that in high glucose, cyst formation is more variable than epithelial layer formation. However, the variability between the two features is similar in physiological glucose and slightly higher than that in high glucose (Figure 2C). Notably, morphological variability for both glucose conditions increased in the neural induction stage after day 6 compared to the hPSC growth stage (Figure 2D).

**Figure 2.**
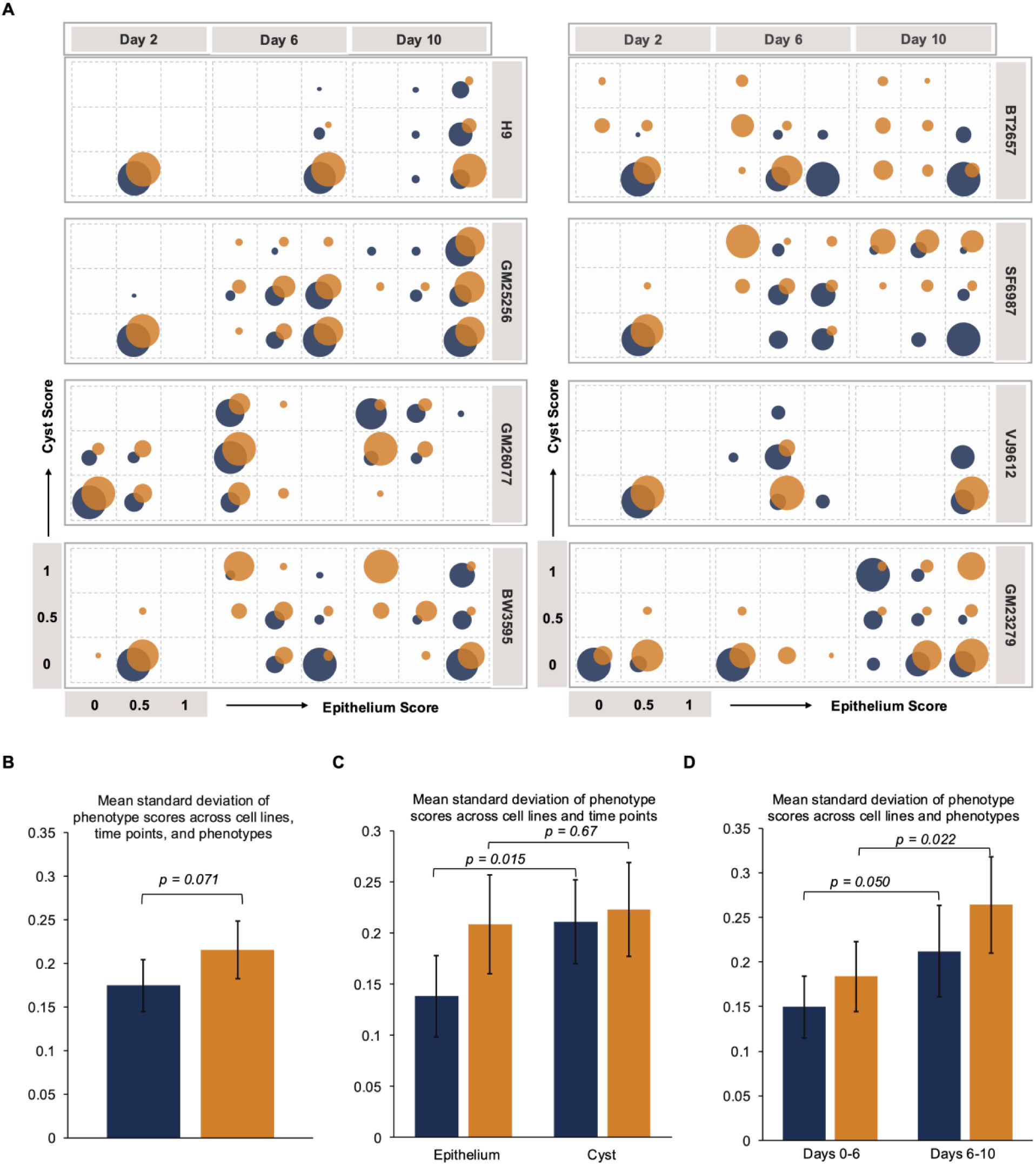
Glucose conditions and growth stage influence morphological heterogeneity in hCOs. A) Frequency bubble plots displaying hCO phenotype score evolution between Day 2 and Day 10 of growth for high (blue) and physiological (orange) glucose conditions for different cell lines. Three levels of epithelium (x-axis) and cyst (y-axis) phenotype scores were assigned, 0 – absence of feature, 0.5 – partially visible / partially developed feature, 1 – clearly visible feature (Supplementary Figure S2). B-D) The mean standard deviation of phenotype scores comparing high (blue) vs physiological (orange) glucose. Standard deviations were calculated independently for each phenotype, time point, and cell line and means of those standard deviations were calculated for the indicated subsets of conditions (Supplementary Figures S3-S4). All error bars represent 95% confidence intervals. All conditions were compared using a two-sample t-test assuming unequal variances. The numbers of hCOs measured are in Supplementary Table S3.

### 3.3 Glucose consumption increases in the neural induction stage

We next asked how glucose concentrations might be driving hCO phenotypes. Potential osmotic effects were controlled for through the use of L-glucose to maintain equivalent osmotic pressure in all conditions. Therefore, we hypothesized that differences in glucose consumption rates between cell lines contribute to the observed morphological variation. Additionally, there is a possibility that hCOs might be completely deprived of glucose between media changes if their consumption rates are high enough. To address this, we selected 4 cell lines that captured different derivation methods as well as phenotypic responses (Supplementary Table S1), generated hCOs, and measured their glucose consumption over time. At each media change, spent media was collected, and their glucose levels were measured. Glucose consumption over time exhibited some common as well as cell line specific patterns (Figures 3A). We observed that even under physiological glucose conditions, hCOs were never entirely deprived of glucose, with at least ∼1 mM remaining at all times. In high glucose conditions, hCOs consumed more total glucose (Figure 3B); however, the fraction of available glucose consumed was higher under physiological glucose conditions (Figure 3C). Furthermore, glucose consumption increased during the neural induction stage compared to the hPSC growth stage, under both high and physiological glucose conditions, likely due to the presence of more cells at this stage (Figure 3B-C).

**Figure 3.**
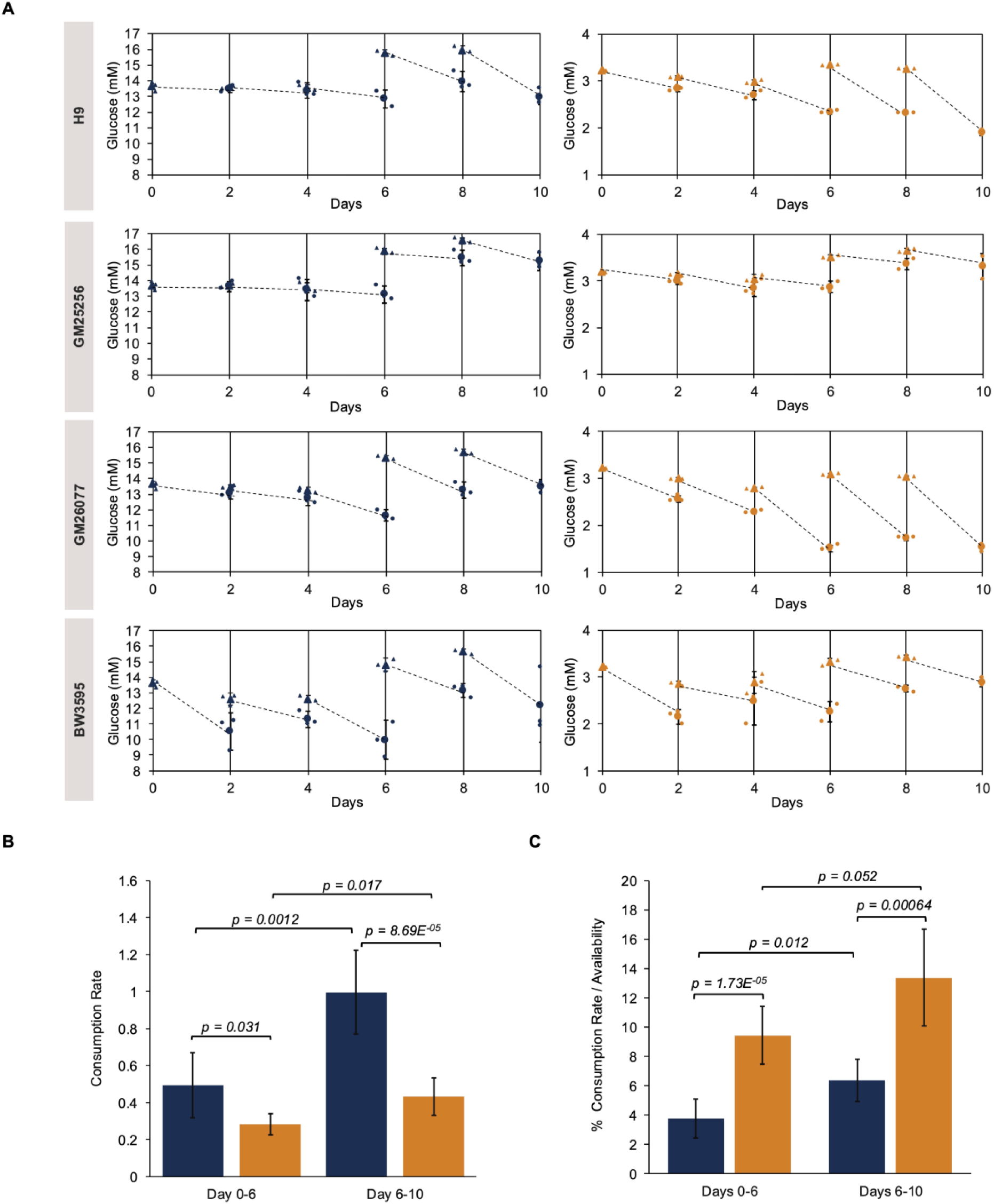
Glucose consumption increases in the neural induction stage. A) Change in media glucose concentration over time in high (blue) vs physiological (orange) glucose conditions. Triangles represent glucose concentration available to hCOs after partial media change. Circles represent glucose concentration in spent media. Dotted lines indicate the drop in mean glucose concentration between media changes. B) Mean glucose consumption rate (change in concentration (mM) / time (days)) and C) Mean glucose consumption rate / initial glucose availability (n = 38 hCOs) across cell lines in high (blue) vs physiological (orange) glucose conditions in hPSC growth stage (Day 0-6) and neural induction stage (Day 6-10). All error bars represent 95% confidence intervals. All conditions were compared using a two-sample t-test assuming unequal variances.

### 3.4 Physiological glucose conditions result in the loss of neurodevelopmental cell types in a cell line–dependent manner

Alterations in cellular metabolism, exhibited here by variable glucose consumption, are known to impact differentiation patterns in hCO models (33). Therefore, to investigate potential variations in differentiation and cell-type composition, we quantified fractions of common neurodevelopmental cell types in 11 days old hCOs: SOX2+ neural precursors, PAX6+ neural ectodermal cells, phospho-vimentin+ mitotic radial glia, and TUJ1+ neurons. Averaging all cell lines, the fraction of SOX2+, PAX6+, and TUJ1+ neurons was not affected by physiological glucose (Figure 4A). However, we also found that hCOs generated from the 4 iPSCs reprogrammed from blood cells (BT2657, BW3595, SF6987, VJ9612) exhibited lower fractions of TUJ1 than all other lines in physiological glucose (Figure 4B). In these blood-derived cell lines, physiological glucose conditions lead to fewer fractions of neurons, with 14 out of 22 hCOs showing no TUJ1+ expression compared to high glucose conditions where only 2 of 21 hCOs showed no TUJ1+ expression (Figures 4C-D). For BT2657, the SOX2+ neural precursor subpopulation was also sparse in physiological glucose conditions (Supplementary Figure S6B). In two cell lines (GM25256 and H9), an intermediate neurodevelopmental cell type: PAX6+ neuroectodermal cells were found to be significantly reduced (Figures 4E, Supplementary Figure S7). However, on the contrary, expression of an earlier developmental cell type, SOX2+ neural precursors, remained unchanged between glucose conditions (Figure 4F, Supplementary Figure S6B). Notably, GM26077 iPSC line showed lower SOX2 expression, at least ∼60% lower than other cell lines, in both high and physiological glucose conditions (Supplementary Figure S6A-B). Emergence of phospho-vimentin+ mitotic radial glia was unaffected by glucose availability and consumption across cell lines, with the exception of GM25256 (Supplementary Figure S8). These results suggest that physiological glucose conditions might be suppressing differentiation at an intermediate stage between neural precursors and post-mitotic neurons, and underscore the potential influence of genotype, donor cell type prior to reprogramming, and glucose levels on varying cell type compositions and developmental trajectories of hCOs.

**Figure 4.**
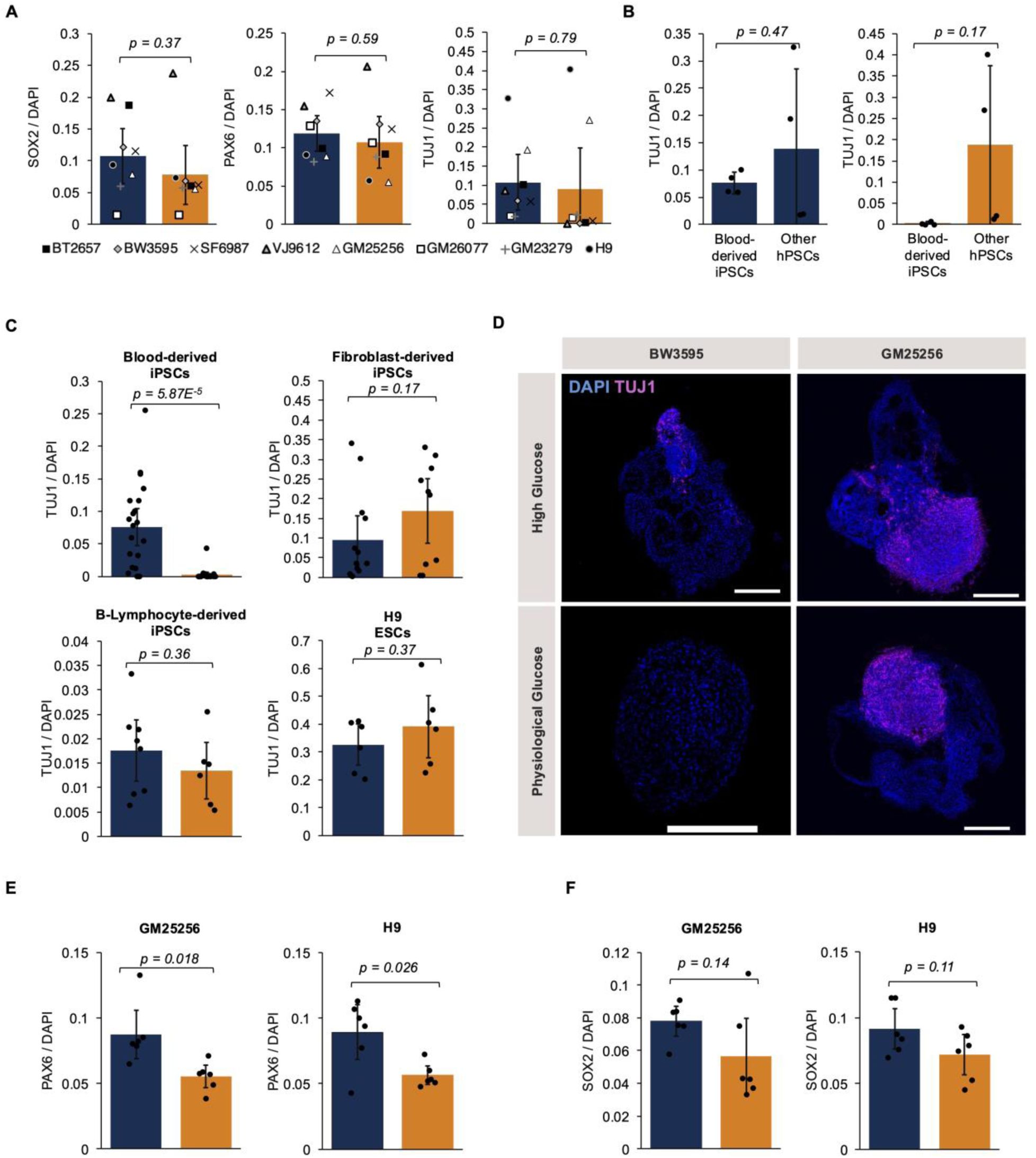
Physiological glucose conditions result in the loss of neurodevelopmental cell types in a cell line–dependent manner. A) Mean fraction of SOX2+ neural precursor cells, PAX6+ neuroectodermal cells, and TUJ1+ neurons relative to DAPI across all cell lines in high (blue) vs physiological (orange) conditions. Each marker represents the mean fraction within each cell line and error bars represent 95% confidence intervals. B) Mean fraction of TUJ1+ neurons relative to DAPI in blood-derived iPSCs compared to the other PSCs in high glucose and physiological glucose conditions. Each black dot represents an individual cell line and error bars represent 95% confidence intervals. C) Mean fraction of TUJ1+ neurons relative to DAPI (nuclei) in hCOs derived from different pluripotent cell lines grouped by their source, comparing high (blue) vs physiological (orange) glucose conditions. D) Representative day 10 hCO images from blood (BW3595) and fibroblast (GM25256) derived iPSCs showing DAPI (nuclei-blue) and TUJ1 (neurons-magenta) in high vs physiological glucose conditions. All scale bars are 250 μm. E) Mean fraction of PAX6+ neuroectodermal cells and F) SOX2+ neural precursors relative to DAPI (nuclei) in hCOs derived from GM25256 and H9 PSCs comparing high (blue) vs physiological (orange) glucose conditions. All fraction quantifications are based on immunofluorescence images. All black dots represent individual hCOs (C, E-F). The number of hCOs measured are in Supplementary Table S4. All conditions were compared using a two-sample t-test assuming unequal variances.

### 3.5 Rapamycin treatment partially recapitulates physiological glucose phenotypes

Glucose deficiency has been linked to the inhibition of the mammalian target of rapamycin (mTOR) cellular signaling pathway (34). To investigate the role of this pathway in glucose-driven morphological and cell-type composition changes, we inhibit mTOR using rapamycin in H9-ESC derived hCOs. H9 hCOs were chosen for this experiment due to their greatest batch-to-batch consistency in growth and morphology in our hands. Over 10 days, hCOs exposed to rapamycin showed slower growth compared to hCOs exposed to vehicle control (DMSO), in both high and physiological glucose conditions (Figure 5A-B), consistent with previous studies showing a negative effect of rapamycin on cell growth (35). Interestingly, in the presence of rapamycin, physiological glucose conditions produced larger hCOs compared to the high glucose conditions, opposite to the trend observed in vehicle control (Figure 5A-B). Additionally, the tendency of hCOs in high glucose to develop cyst-like features was reduced in the presence of rapamycin (Figure 5C, Supplementary Figure S9A), making it comparable to the morphology under physiological glucose conditions. Unexpectedly, immunofluorescence analysis demonstrated that the loss of SOX2+ neural precursors and Ki67+ proliferative cells observed in physiological glucose conditions was rescued by the addition of rapamycin (Figure 5D-E). Conversely, the fraction of TUJ1+ neurons reduced under physiological glucose with rapamycin treatment (Figure 5F). These results suggest mTOR signaling and glucose levels may have a complex relationship in regulating hCO development and differentiation.

**Figure 5.**
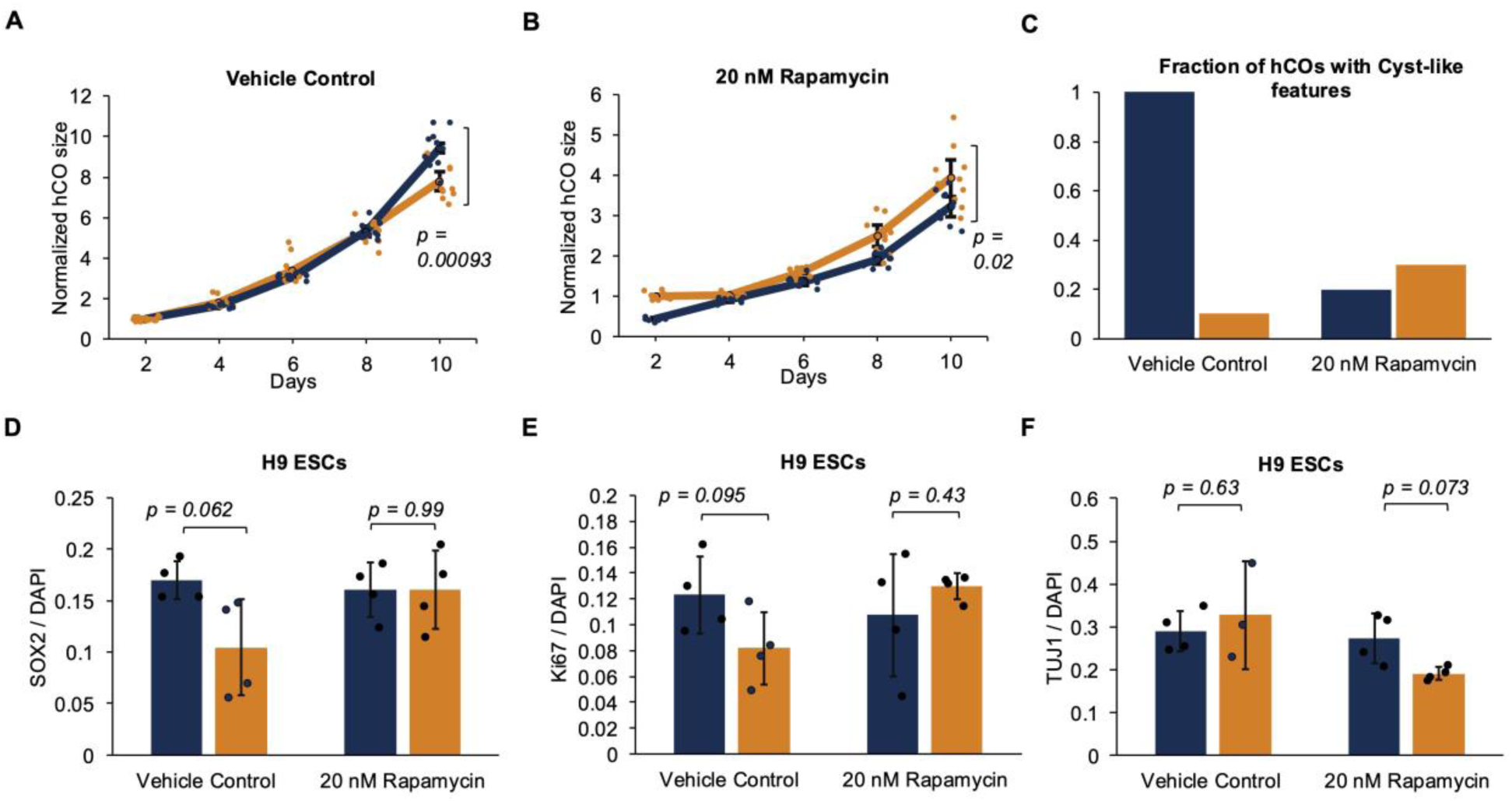
Rapamycin treatment partially recapitulates physiological glucose phenotypes. H9 ESC-derived hCO growth over 10 days in high (blue) and physiological glucose (orange) conditions for hCOs exposed to A) vehicle control (DMSO) and B) 20 nM Rapamycin. hCO sizes on the y-axis are normalized to the corresponding average size of physiological glucose hCOs on Day 2. Solid lines connect the average hCO size between time points and dots represent individual hCO sizes. hCO sizes at Day 10 were compared using a two-sample t-test assuming unequal variances. C) Fraction of hCOs with cyst-like phenotypes at 10 days in high (blue) vs physiological (orange) glucose upon exposure to vehicle control (DMSO) and 20 nM Rapamycin, calculated based on Supplementary Figure 9A. Mean fraction of D) SOX2+ neural precursors, E) proliferating Ki67+ cells, and F) TUJ1+ neurons relative to DAPI (nuclei) in hCOs in high (blue) vs physiological (orange) glucose conditions upon exposure to vehicle control (DMSO) and 20 nM Rapamycin. All marker quantifications are based on immunofluorescence images. All black dots represent individual hCOs and all error bars represent 95% confidence intervals. The numbers of hCOs measured can be found in Supplementary Table S5. All conditions were compared using a two-sample t-test assuming unequal variances.

## 4. Discussion

We examined whole brain hCOs derived from eight human pluripotent stem cell lines and tracked glucose-induced changes in growth, morphology, and cell-type composition over 10 days. We observe that differences between high and physiological glucose conditions were more pronounced beyond day 6 across cell lines. Since the culture conditions are switched from a proliferation supporting medium to a neural fate induction medium at day 6, it is possible that the new media components interact with glucose differently to drive a greater change in phenotype, particularly morphology. Simultaneously, by day 6 hCOs are larger, and glucose consumption rates would therefore be expected to increase. So, it might be that physiological glucose conditions require more frequent media changes to maintain a minimal level for productive hCO development. Indeed, we observed more substantial depletion of glucose during the latter periods of our experiments (days 6-10), Figure 3A). While there is still some basal level of glucose present at all times, it may be that glucose concentrations are required to be above a minimal level for cells to effectively consume it.

Here, we focused on two prominent morphological features – neuroepithelium and cysts – which have both been previously noted in hCO cultures. While neuroepithelium is deemed as a desired or expected feature for hCOs, tissues with cyst-like features are considered undesirable or failed hCOs (29,36). GM26077 and GM23279 iPSC-derived hCOs showed limited neuroepithelium formation but cyst-like features emerged even in typical, high glucose conditions suggesting the stem cells may not be suitably primed for hCO cultures. A recent study showed that presence of cysts features can also indicate direction towards mesenchymal fate instead of neural fate (37). The role of glucose in driving this cyst-like phenotype remains unclear, but physiological glucose condition leads to loss of neurodevelopmental cell types in multiple cell lines. Some cell lines such as BT2657, BW3595, and SF6987 showed more than 40% increases in cyst-like features and a lack of TUJ1+ neurons in physiological glucose conditions. However, conversely, H9- derived ESCs showed a ∼70% decrease in cyst-like features in physiological glucose conditions, accompanied with no loss of TUJ1+ neurons; it did show significant reduction in PAX6+ neural ectodermal cell populations. Interestingly, prior studies investigating murine neural stem cells show that impaired proliferation (38) and differentiation to neurons (15,39,40), and neural tube formation defects (41), are observed in high glucose (∼25 mM) conditions compared to physiological (∼2-5 mM) glucose conditions. Human cell-based models of neurodevelopment have also shown inhibition of differentiation at hyperglycemic glucose concentrations > 25 mM (23,25,42). These mixed findings suggest hCO development may be non-linearly affected by glucose levels, and exhibit variation based upon the source and derivation of each cell line.

Previous work has considered the role of glucose in *in vitro* neurodevelopment in combination with other cell culture media constituents. Prior research with human neural stem cells demonstrated that lowering both glucose and insulin to physiological concentrations supports neurosphere formation and differentiation to cortical fate (43). Bardy and colleagues developed a novel Brain-Phys medium for human iPSC-derived neurons and showed that physiological media conditions improve electrophysiological properties (16). Based on these investigations, we speculate that while reducing glucose alone may impair neural fate induction and differentiation, concurrently modulating other interacting metabolites (e.g., amino acids, lipids) and signaling molecules (e.g., insulin) could improve hCO models. Moreover, non-physiological conditions in hCOs have been shown to mask disease phenotypes *in vitro*, that can be rescued under physiological conditions (44).

Further investigations could also elucidate the mechanism behind glucose driving neurodevelopmental changes in hCO models. Glucose metabolism is linked to mTOR (mammalian target of rapamycin) signaling, which is a regulator of cell survival, growth, and proliferation (45,46). Under glucose deprivation conditions, increased AMP/ATP ratios are sensed by AMPK leading to inhibition of mTOR complex 1 (mTORC1) (34,47) which leads to suppression of cell proliferation (46). To evaluate the mechanistic role of mTOR in glucose-driven neural differentiation, we exposed hCOs in both high and physiological glucose conditions to 20 nM rapamycin, an mTOR inhibitor. In high glucose conditions, the neurodevelopmental marker expression was not downregulated in the presence of rapamycin. It is possible that while fractions of cell types remain unchanged, cellular morphologies and architecture might be affected by rapamycin (48,49). We did observe an expected reduction in hCO size but unexpectedly observed larger hCOs in physiological compared to high glucose, suggesting potential synergistic or antagonistic interactions between lower glucose levels and mTOR inhibition (50).

This study underscores that hCO growth, morphology, and cell-type composition can differ widely between cell lines. It also draws attention to the role glucose plays as a morphogen in hCO models. The findings point to the importance of exploring additional metabolic components in the culture media, as these may offer new opportunities for enhancing hCO models and for unraveling how metabolism intersects with signaling pathways.

## Acknowledgements

We thank Christopher Cummings, Suleiman Sweilem, and Laurie Overton at the Golden LEAF Biomanufacturing Training and Education Center (BTEC), North Carolina State University for access to the Cedex^®^ Bio Analyzer (Roche Diagnostics) and performing the glucose measurements on spent media. We thank Mahe Jabeen and John Britt for valuable insights on cellular metabolism. We thank Dr. Maria Theresa Fadri, Z. Begum Yagci, Leandra Caywood, Tyler J. Johnson, and Rachel Polak for helpful discussions and feedback.

## 6. Author Contributions

Conceptualization: GRK, BMR, AJK.

Methodology: GRK.

Investigation: GRK, AJK.

Visualization: GRK.

Supervision: AJK.

Writing - Original draft: GRK, AJK.

Writing – Review and Editing: GRK, BMR, AJK.

## 9. Data Availability

The datasets used to generate the graphs in this study can be found in the supplementary materials.

