## Supplementary Material for "Glucose levels impact the morphology and cell-type composition of human cerebral organoids"

### Supplementary Figures

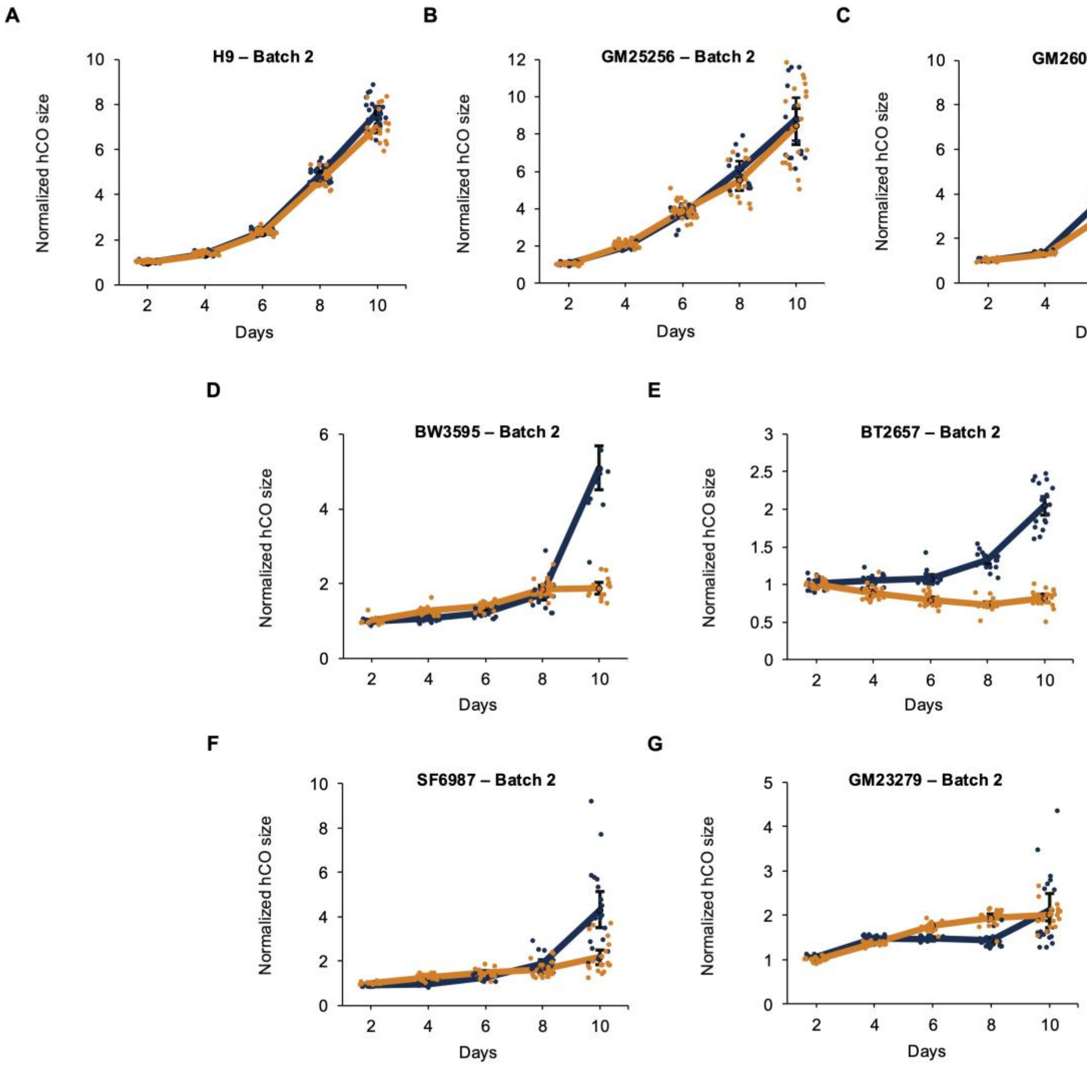


**Supplementary Figure S1.** hCO growth over 10 days in high (blue) and physiological glucose (orange) conditions for hCOs generated from different cell lines. hCO sizes on the y-axis are normalized to the corresponding average size of physiological glucose hCOs on Day 2. Solid lines connect the mean hCO size between time points, dots represent individual hCO sizes, and error bars represent 95% confidence intervals. The numbers of hCOs measured can be found in Supplementary Table S2.

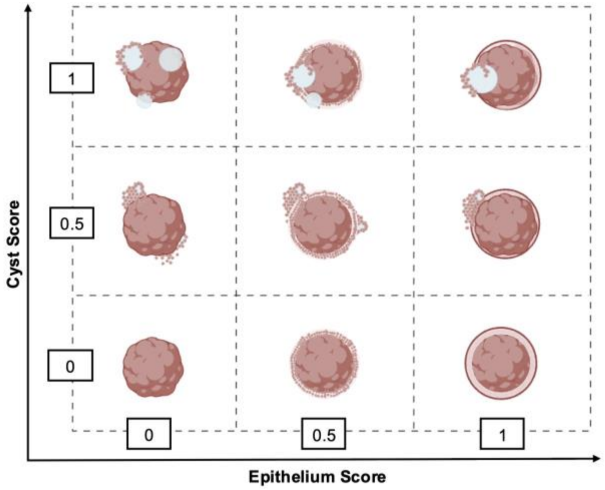


**Supplementary Figure S2.** hCO scoring scheme. Three levels of epithelium (x-axis) and cyst (y-axis) phenotype scores were assigned, 0 – absence of feature, 0.5 – partially visible / partially developed feature, 1 – clearly visible feature.

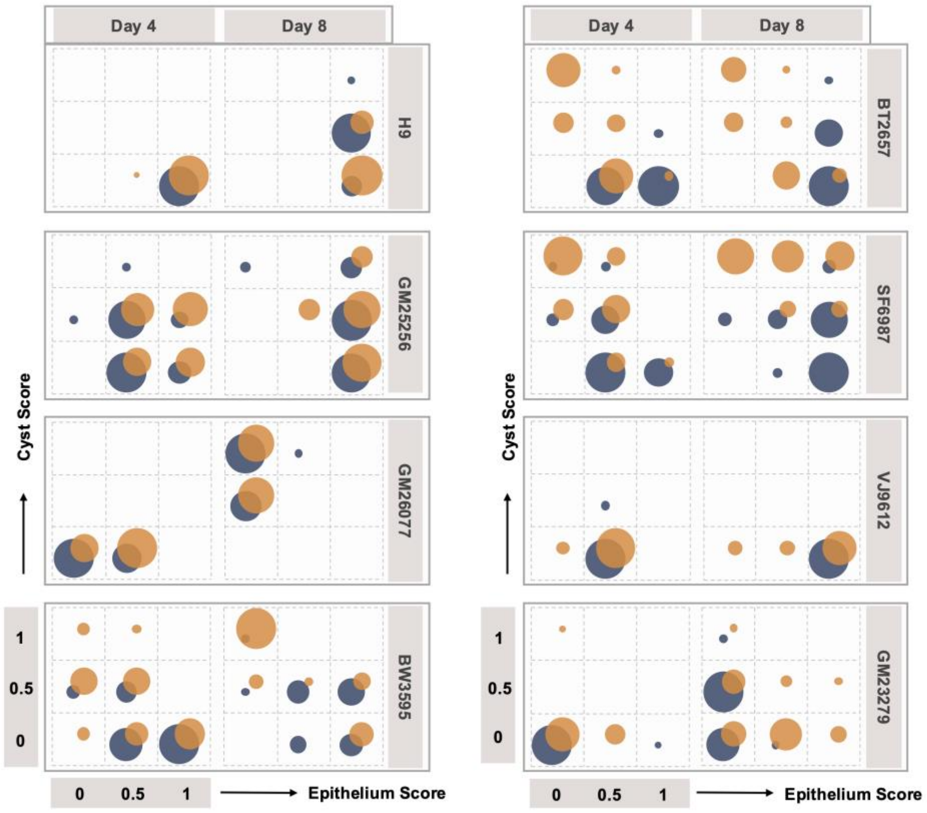


**Supplementary Figure S3.** Frequency bubble plots displaying hCO phenotype scores on days 4 and 8 for high (blue) and physiological (orange) glucose conditions for different cell lines. The hCO scoring scheme in Supplementary Figure S2 was followed to generate these plots. The numbers of hCOs measured are in Supplementary Table S3.

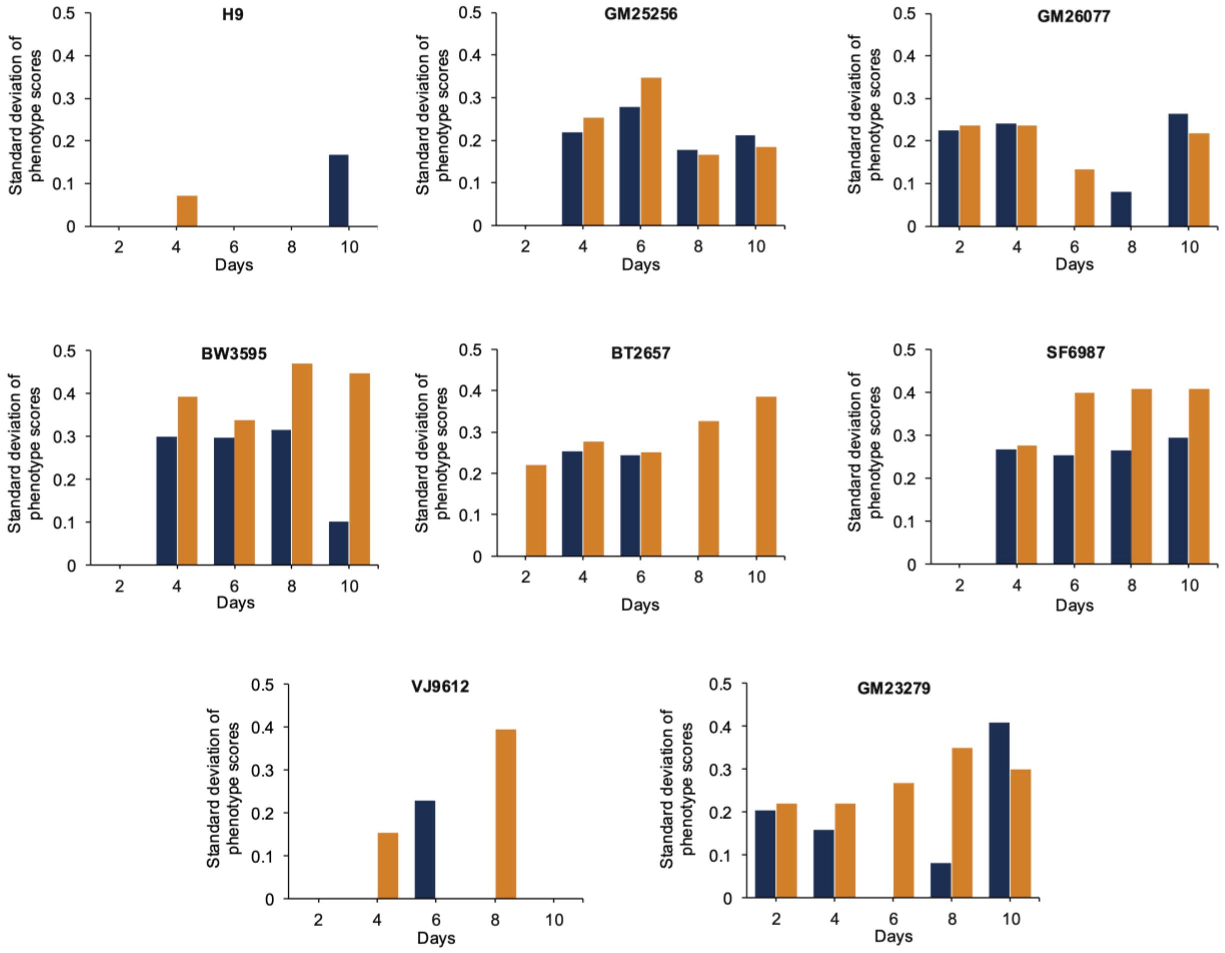


**Supplementary Figure S4.** The standard deviation of hCO phenotype scores comparing high (blue) vs physiological (orange) glucose conditions for the epithelium phenotype at each time for all cell lines.

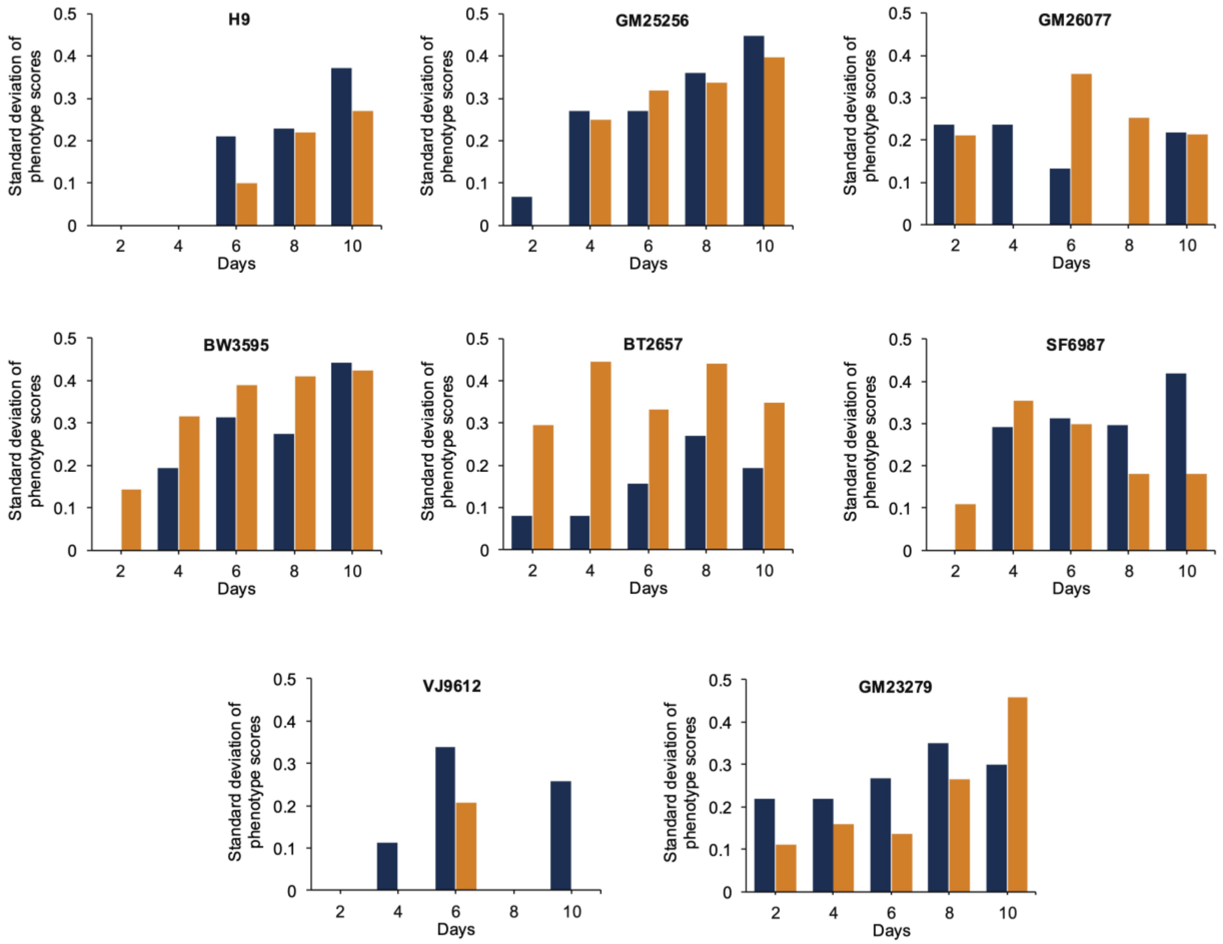


**Supplementary Figure S5.** The standard deviation of hCO phenotype scores comparing high (blue) vs physiological (orange) glucose conditions for the cyst phenotype at each time for all cell lines.

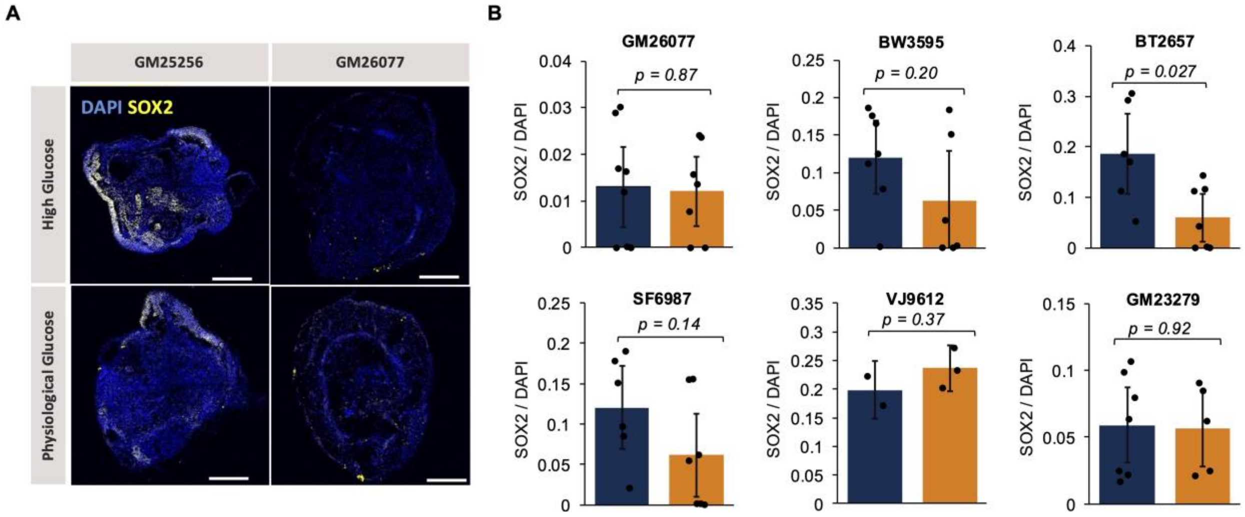


**Supplementary Figure S6.** A) Immunofluorescence images showing SOX2 (neural precursors-yellow) and DAPI (nuclei-blue) expression in GM25256 and GM26077-derived hCOs at 11 days in high vs physiological glucose. All scale bars are 250 μm. B) Mean fraction of SOX2+ neural precursors relative to DAPI in hCOs derived from different cell lines for high (blue) vs physiological (orange) glucose conditions. All fraction quantifications are based on immunofluorescence images. All black dots represent individual hCOs and error bars represent 95% confidence intervals. The numbers of hCOs measured are in Supplementary Table S4. All conditions were compared using a two-sample t-test assuming unequal variances.

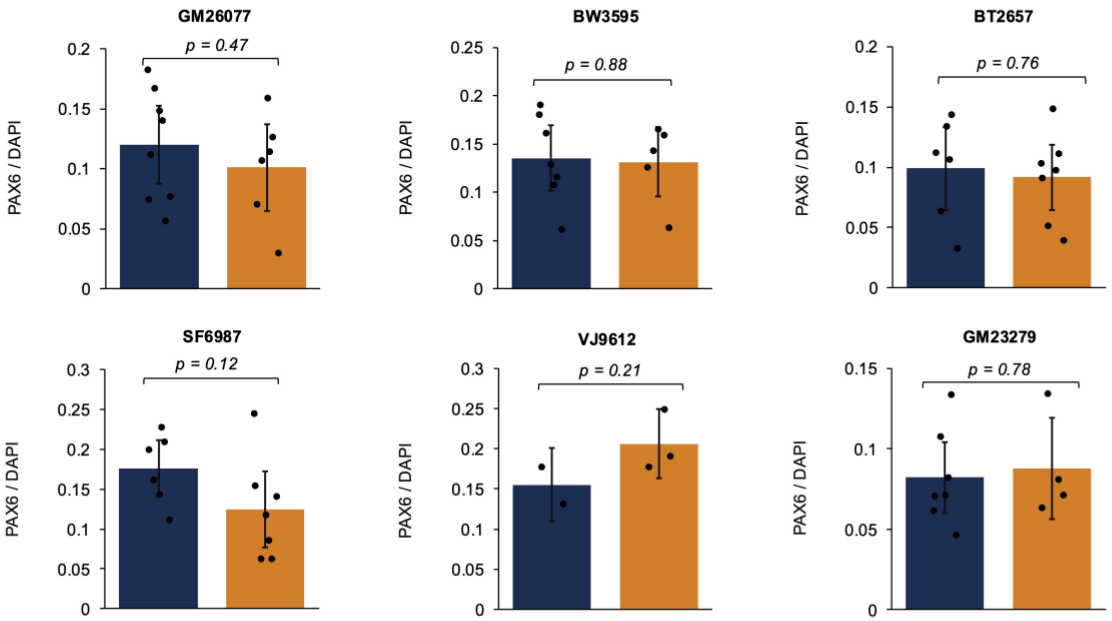


**Supplementary Figure S7.** Mean fraction of PAX6+ neuroectodermal cells relative to DAPI (nuclei) in hCOs derived from different cell lines for high (blue) vs physiological (orange) glucose conditions. All fraction quantifications are based on immunofluorescence images. All black dots represent individual hCOs and error bars represent 95% confidence intervals. The numbers of hCOs measured are in Supplementary Table S4. All conditions were compared using a two-sample t-test assuming unequal variances.

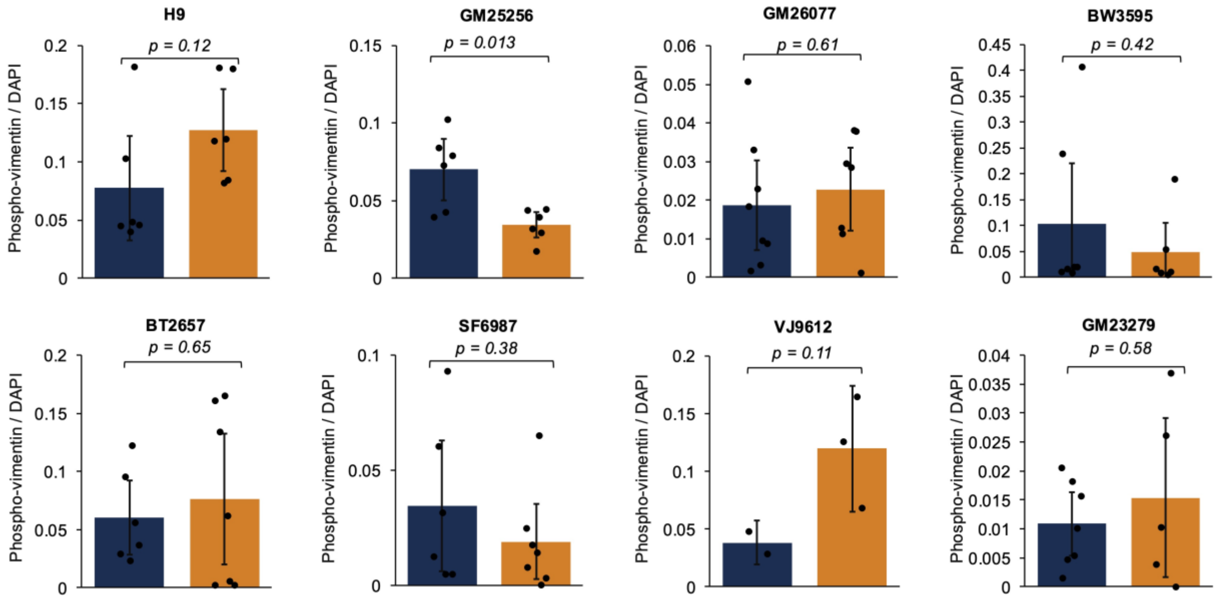


**Supplementary Figure S8.** Mean fraction of phospho-vimentin+ mitotic radial glia relative to DAPI (nuclei) in hCOs derived from different cell lines for high (blue) vs physiological (orange) glucose conditions. All fraction quantifications are based on immunofluorescence images. All black dots represent individual hCOs and error bars represent 95% confidence intervals. The numbers of hCOs measured are in Supplementary Table S4. All conditions were compared using a two-sample t-test assuming unequal variances.

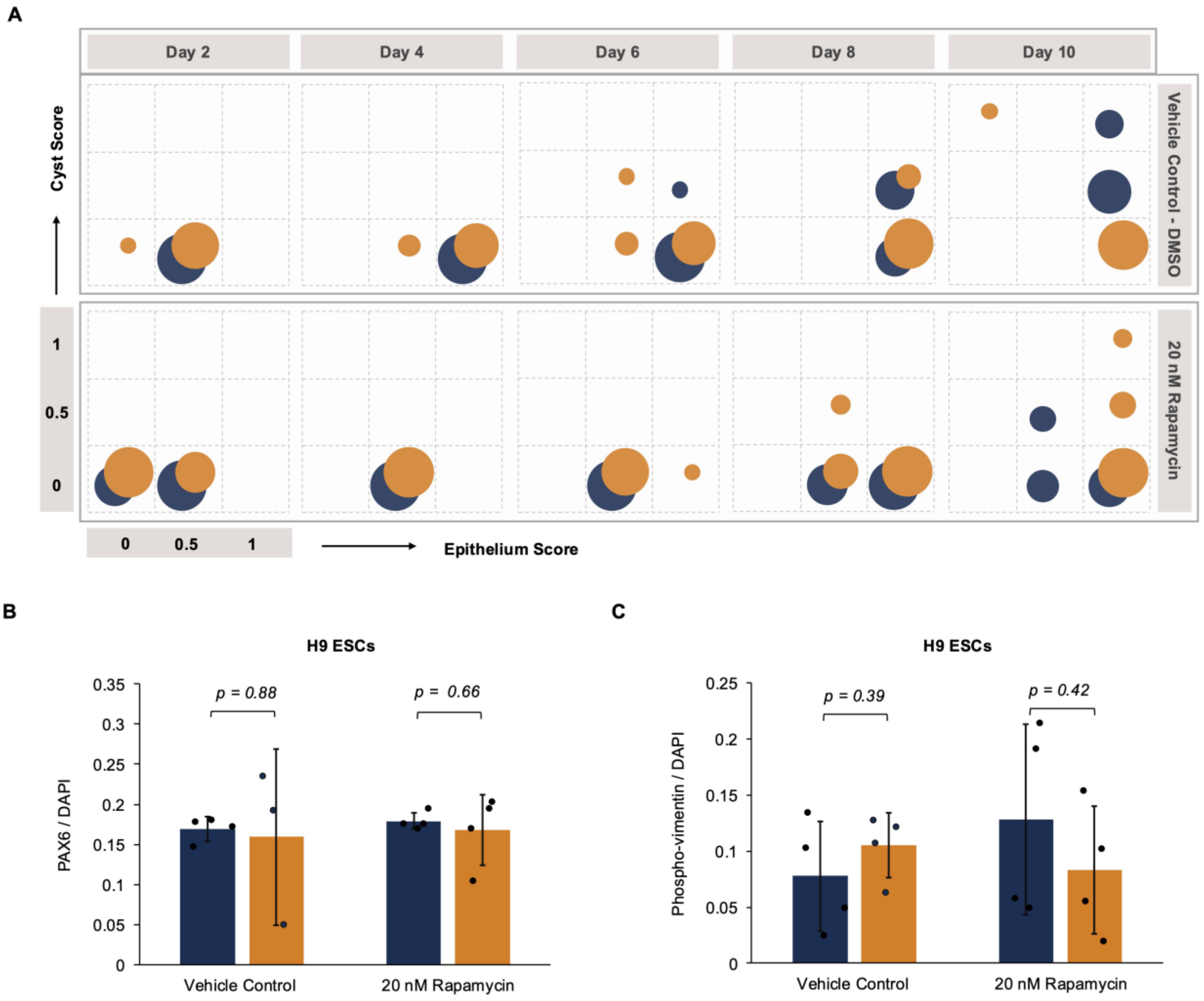


**Supplementary Figure S9.** A) Frequency bubble plots displaying hCO phenotype score evolution between Day 2 and Day 10 of growth for high (blue) and physiological (orange) glucose conditions for for H9 ESC-derived hCOs exposed to the vehicle control (DMSO) and 20 nM Rapamycin (n=10 hCOs measured per subplot per condition). The hCO scoring scheme in Supplementary Figure S2 was followed to generate these plots. Mean fraction of B) PAX6+ neuroectodermal cells and C) phospho-vimentin+ mitotic radial glia relative to DAPI (nuclei) in hCOs in high (blue) vs physiological (orange) glucose conditions upon exposure to vehicle control (DMSO) and 20 nM Rapamycin. All marker quantifications are based on immunofluorescence images. All black dots represent individual hCOs and all error bars represent 95% confidence intervals. The numbers of hCOs measured can be found in Supplementary Table S5. All conditions were compared using a two-sample t-test assuming unequal variances.

**Supplementary Tables**

**Supplementary Table S1: Information on the hPSC cell lines.**

| Cell line | Type | Sex | Source |
|---|---|---|---|
| H9 | ESC | F | -/- |
| SF6987 | iPSC (Gift from Yale University) | F | Blood |
| BT2657 | iPSC (Gift from Yale University) | F | Blood |
| BW3595 | iPSC (Gift from Yale University) | F | Blood |
| VJ9612 | iPSC (Gift from Yale University) | M | Blood |
| GM26077 | iPSC (Coriell Institute for Medical Research) | F | B-Lymphocyte |
| GM23279 | iPSC (Coriell Institute for Medical Research) | F | Fibroblasts |
| GM25256 | iPSC (Coriell Institute for Medical Research) | M | Fibroblasts |

**Supplementary Table S2: The number of hCOs measured for size analysis at different time points comparing high vs physiological glucose conditions for all cell lines.**

| Sub-Figure | High Glucose | Physiological Glucose |
|---|---|---|
| Figure 1B - H9 | 20 | 20 |
| Figure 1B - GM25256 | 18-24 | 16-23 |
| Figure 1B - GM26077 | 19-20 | 20 |

| | | |
|---|---|---|
| Figure 1B – BW3595 | 10-20 | 18-20 |
| Figure 1B – BT2657 | 19-20 | 14-20 |
| Figure 1B – SF6987 | 17-20 | 18-20 |
| Figure 1B – VJ9612 | 7-20 | 7-20 |
| Figure 1B – GM23279 | 20 | 19-20 |
| Supplementary Figure S1 - H9 | 20-30 | 20-30 |
| Supplementary Figure S1 - GM25256 | 15-30 | 15-30 |
| Supplementary Figure S1 - GM26077 | 20 | 20 |
| Supplementary Figure S1 – BW3595 | 14-20 | 14-20 |
| Supplementary Figure S1 – BT2657 | 18-19 | 20 |
| Supplementary Figure S1 – SF6987 | 20 | 20 |
| Supplementary Figure S1 – GM23279 | 19-20 | 19-20 |

**Supplementary Table S3: The number of hCOs measured for phenotype analysis in Figure 2 and Supplementary Figure S3 at different time points comparing high vs physiological glucose conditions for all cell lines.**

| Cell Line | High Glucose | | | | | Physiological Glucose | | | | |
|---|---|---|---|---|---|---|---|---|---|---|
| | Day 2 | Day 4 | Day 6 | Day 8 | Day 10 | Day 2 | Day 4 | Day 6 | Day 8 | Day 10 |
| H9 | 50 | 49 | 50 | 38 | 40 | 50 | 50 | 50 | 40 | 40 |
| GM25256 | 54 | 54 | 54 | 32 | 36 | 53 | 54 | 54 | 33 | 36 |
| GM26077 | 40 | 40 | 40 | 39 | 39 | 40 | 40 | 40 | 40 | 40 |

| | | | | | | | | | | |
|---|---|---|---|---|---|---|---|---|---|---|
| BW3595 | 40 | 39 | 40 | 30 | 24 | 40 | 40 | 40 | 38 | 32 |
| BT2657 | 39 | 39 | 38 | 38 | 39 | 40 | 40 | 34 | 34 | 36 |
| SF6987 | 40 | 40 | 37 | 40 | 37 | 40 | 40 | 38 | 40 | 40 |
| VJ9612 | 20 | 20 | 15 | 7 | 16 | 20 | 20 | 15 | 7 | 16 |
| GM23279 | 40 | 40 | 39 | 39 | 39 | 40 | 40 | 38 | 39 | 39 |

**Supplementary Table S4: The number of hCOs measured for neurodevelopmental marker quantification in high vs physiological glucose conditions for all cell lines.**

| Sub-Figure | High Glucose | Physiological Glucose |
|---|---|---|
| Figure 1C – H9 (1-2 images per hCO) | 6 | 6 |
| Figure 1C – GM25256 (1-2 images per hCO) | 6 | 6 |
| Figure 1C – GM26077 (1-2 images per hCO) | 8 | 7 |
| Figure 1C – BW3595 (1 image per hCO) | 7 | 6 |
| Figure 1C – BT2657 (1 image per hCO) | 6 | 7 |
| Figure 1C – SF6987 (1-2 images per hCO) | 6 | 7 |
| Figure 1C – VJ9612 (1 image per hCO) | 2 | 3 |
| Figure 1C – GM23279 (1 image per hCO) | 7 | 5 |
| Figure 4C – Blood-derived (1-2 images per hCO) | 21 | 22 |
| Figure 4C – Fibroblast-derived (1-2 images per hCO) | 13 | 10 |
| Figure 4C B-Lymphocyte-derived (1-2 images per hCO) | 8 | 6 |
| Figure 4C H9 (1-2 images per hCO) | 6 | 6 |
| Figure 4E – GM25256 (1-2 images per hCO) | 6 | 6 |

| | | |
|---|---|---|
| Figure 4E – H9 (2-3 images per hCO) | 6 | 6 |
| Figure 4F – GM25256 (4-6 images per hCO) | 6 | 6 |
| Figure 4F – H9 (4-6 images per hCO) | 6 | 6 |
| Supplementary Figure S6 – GM26077 (1-4 images per hCO) | 8 | 7 |
| Supplementary Figure S6 – BW3595 (1-2 images per hCO) | 7 | 6 |
| Supplementary Figure S6 – BT2657 (2 images per hCO) | 6 | 7 |
| Supplementary Figure S6 – SF6987 (2-4 images per hCO) | 6 | 7 |
| Supplementary Figure S6 – VJ9612 (2 images per hCO) | 2 | 3 |
| Supplementary Figure S6 – GM23279 (1-3 images per hCO) | 7 | 5 |
| Supplementary Figure S7 – GM26077 (1-2 images per hCO) | 8 | 6 |
| Supplementary Figure S7 – BW3595 (1 image per hCO) | 7 | 5 |
| Supplementary Figure S7 – BT2657 (1 image per hCO) | 6 | 7 |
| Supplementary Figure S7 – SF6987 (1-2 images per hCO) | 6 | 7 |
| Supplementary Figure S7 – VJ9612 (1 image per hCO) | 2 | 3 |
| Supplementary Figure S7 – GM23279 (1-2 images per hCO) | 7 | 4 |
| Supplementary Figure S8 – H9 (2-3 images per hCO) | 6 | 6 |
| Supplementary Figure S8 – GM25256 (2-3 images per hCO) | 6 | 6 |
| Supplementary Figure S8 – GM26077 (1-2 images per hCO) | 8 | 7 |
| Supplementary Figure S8 – BW3595 (1 image per hCO) | 7 | 6 |
| Supplementary Figure S8 – BT2657 (1 image per hCO) | 6 | 7 |
| Supplementary Figure S8 – SF6987 (1-2 images per hCO) | 6 | 7 |
| Supplementary Figure S8 – VJ9612 (1 image per hCO) | 2 | 3 |
| Supplementary Figure S8 – GM23279 (1 image per hCO) | 7 | 5 |

**Supplementary Table S5: Inhibitor treatment experiment. The number of hCOs measured for size analysis and neurodevelopmental marker quantification comparing high vs physiological glucose conditions for all cell lines.**

| Sub-Figure | High Glucose – 20 nM Rapamycin | Physiological Glucose – 20 nM Rapamycin | High Glucose - Vehicle Control | Physiological Glucose – Vehicle Control |
|---|---|---|---|---|
| Figure 5A | -/- | -/- | 10 | 10 |
| Figure 5B | 10 | 10 | -/- | -/- |
| Figure 5D (2-6 images per hCO) | 4 | 4 | 4 | 4 |
| Figure 5E (1-2 images per hCO) | 4 | 4 | 4 | 4 |
| Figure 5F (1-3 images per hCO) | 4 | 4 | 4 | 4 |
| Supplementary Figure S9B (1-3 images per hCO) | 4 | 4 | 4 | 4 |
| Supplementary Figure S9C (1-2 images per hCO) | 4 | 4 | 4 | 4 |
